# Evaluation of p53 Immunohistochemical Expression Using Open-Source Software for Digital Image Analysis. A Tissue Microarray Study of Penile Squamous Cell Carcinomas

**DOI:** 10.1101/029850

**Authors:** Alcides Chaux, Authur L. Burnett, George J. Netto

## Abstract

The addition of molecular biomarkers is needed to increase the accuracy of pathologic factors as prognosticators of outcome in penile squamous cell carcinomas (SCC). Evaluation of these biomarkers is usually carried out by immunohistochemistry. Herein we assess p53 immunohistochemical expression on tissue samples of penile SCC using freely-available, open-source software packages for digital image analysis. We also compared the results of digital analysis with standard visual estimation. Percentages of p53 positive cells were higher by visual estimation than by digital analysis. However, correlation was high between both methods. Our study shows that evaluation of p53 immunohistochemical expression is feasible using open-source software packages for digital image analysis. Although our analysis was limited to penile SCC, the rationale should also hold for other tumor types in which evaluation of p53 immunohistochemical expression is required. This approach would reduce interobserver variability, and would provide a standardized method for reporting the results of immunohistochemical stains. As these diagnostic tools are freely-available online, researchers and practicing pathologists could incorporate them in their daily practice without increasing diagnostic costs.

## Introduction

Most penile tumors are squamous cell carcinomas (SCC) arising at the distal mucosa covering glans, coronal sulcus, or foreskin. Several pathologic features of the primary tumor, including histologic grade, tumor extent, and perineural invasion, have been established as prognosticators of outcome.^1,2^ However, additional markers are needed to increase the accuracy of these pathologic factors. With this goal, immunohistochemical expression of cell cycle-related markers has been used to estimate prognosis of penile SCC.^3^ Nevertheless, differences in evaluation criteria hinder the comparison of series, thwarting the standardization of clinically-useful thresholds of immunohistochemical expression.

Aiming to overcome these difficulties, digital image analysis using proprietary software has been proposed, either for research or clinical practice. However, the high costs of these diagnostic tools preclude their routine implementation. Herein we evaluate open-source software packages, freely available online, to analyze the immunoexpression levels of p53 in penile SCC. We provide universal resource locators (URLs) for downloading these packages, and a basic protocol for digital image analysis. Finally, we also compare the results of digital analysis with standard visual estimation of p53 immunohistochemical expression.

## Material and Methods

### Tissue Microarray Building and Immunohistochemistry

A total of 39 cases of formalin-fixed, paraffin-embedded penile SCC were used to build a tissue microarray (TMA) at the Johns Hopkins TMA Lab Core (Baltimore, MD). Each case was randomly sampled 3–9 times, depending on tumor size, yielding a total of 156 tissue cores of 1 mm of diameter. Pathologic evaluation was done using previously published criteria.^1^

Immunohistochemistry for p53 (antibody against p53, clone-BP53-11, Ventana Medical Systems, Inc. Tucson, AZ) was performed on automated systems from Ventana XT (Ventana Medical Systems, Inc. Tucson, AZ). The reaction was developed using streptavidin-HRP detection I-View kit (Ventana Medical Systems, Inc. Tucson, AZ). All sections were then counterstained with hematoxylin, dehydrated, and cover-slipped.

### Evaluation of p53 Immunohistochemical Expression

Each TMA spot was scanned using the APERIO system (Aperio Technologies, Inc., Vista, CA) and uploaded to TMAJ, an open-source platform for online evaluation of TMA images, available at http://tmaj.pathology.jhmi.edu. Images were scanned at a 20*x* resolution, yielding an image scale of 2.65*μ/mm.* Percentage of p53 positive cells was then established using visual and digital analysis. For this purpose, images were downloaded from the TMAJ database to a local computer.

For visual analysis, percentages of p53 positive nuclei were estimated by naked eye on a computer screen, without the use of any specialized software. For digital analysis, the open-source software ImageJ version 1.44, available at http://rsb.info.nih.gov/ij, was used along with the immunoratio plug-in, available at http://imtmicroscope.uta.fi/immunoratio.

The immunoratio plug-in calculates the percentage of positively stained nuclear area (labeling index) by using a color deconvolution algorithm previously described by Tuominen *et al.*^4^ This deconvolution algorithm separates the staining components (diaminobenzidine and hematoxylin) based on user-defined thresholds for positive nuclei (brown pixels) and negative nuclei (blue pixels). These thresholds were adjusted in a training set of 5 randomly selected TMA spots, until at least 95% of nuclei were identified, either as positive or negative. The same algorithm was then used to estimate in batch the percentage of positive cells. Results were exported afterward to a database containing the pathologic features of the case.

### Statistical Analysis

Analyses were carried out spot by spot and using the pooled arithmetic mean of all the spots for each case. Percentages of p53 positive cells estimated by either visual or digital analyses were compared using the Wilcoxon matched-pairs sign-rank test. The correlation between the visual and the digital methods was evaluated using Spearman’s *ρ* correlation coefficient. Spearman’s *ρ* was interpreted as follows: < 0.09, no correlation; 0.10 − 0.29, weak correlation; 0.30 − 0.49, moderate correlation; ≥ 0.50, strong correlation. The Kruskal-Wallis test was used to compare percentages of p53 stratified by histologic subtype and histologic grade. A 2-tailed *P* < 0.05 was required for statistical significant. Data were analyzed using R version 3.2.2 “Fire Safety”.^5^ The dataset and R scripts used for data analysis, as well as additional results (including the full analysis of the dataset), are freely available at https://github.com/alcideschaux/Penis-p53.

## Results

Table 1 shows the distribution of the 39 cases by histologic subtype and histologic grade. Figure 1 shows scanned TMA spots of H&E and p53-stained tissue cores along with the output of the digital analysis for 1 TMA spot.

**Table 1:** Mean Percentage and Standard Deviation of p53 Positive Cells using Visual Estimation and Digital Analysis, by Histologic Subtype and Grade

|  | No. cases | Visual estimation | P value | Digital estimation | P value |
| --- | --- | --- | --- | --- | --- |
| **Histologic subtype** |  |  | 2.1e-05 |  | 0.0013 |
| Usual SCC | 22 | 31.7 (36.2) |  | 6.4 (9) |  |
| Warty-basaloid SCC | 10 | 8.8 (14.1) |  | 2 (2.7) |  |
| Basaloid SCC | 3 | 0.8 (0.6) |  | 0.3 (0.5) |  |
| Warty SCC | 3 | 42.9 (39.4) |  | 8.2 (9.2) |  |
| Papillary SCC | 1 | 8.3 (2.9) |  | 0.7 (0.6) |  |
| **Histologic grade** |  |  | 0.091 |  | 0.021 |
| Grade 1 | 5 | 14.6 (22.5) |  | 1.8 (2.8) |  |
| Grade 2 | 14 | 35.7 (39.7) |  | 7.4 (10.4) |  |
| Grade 3 | 20 | 13.5 (20.4) |  | 2.9 (3.3) |  |

**Figure 1:**
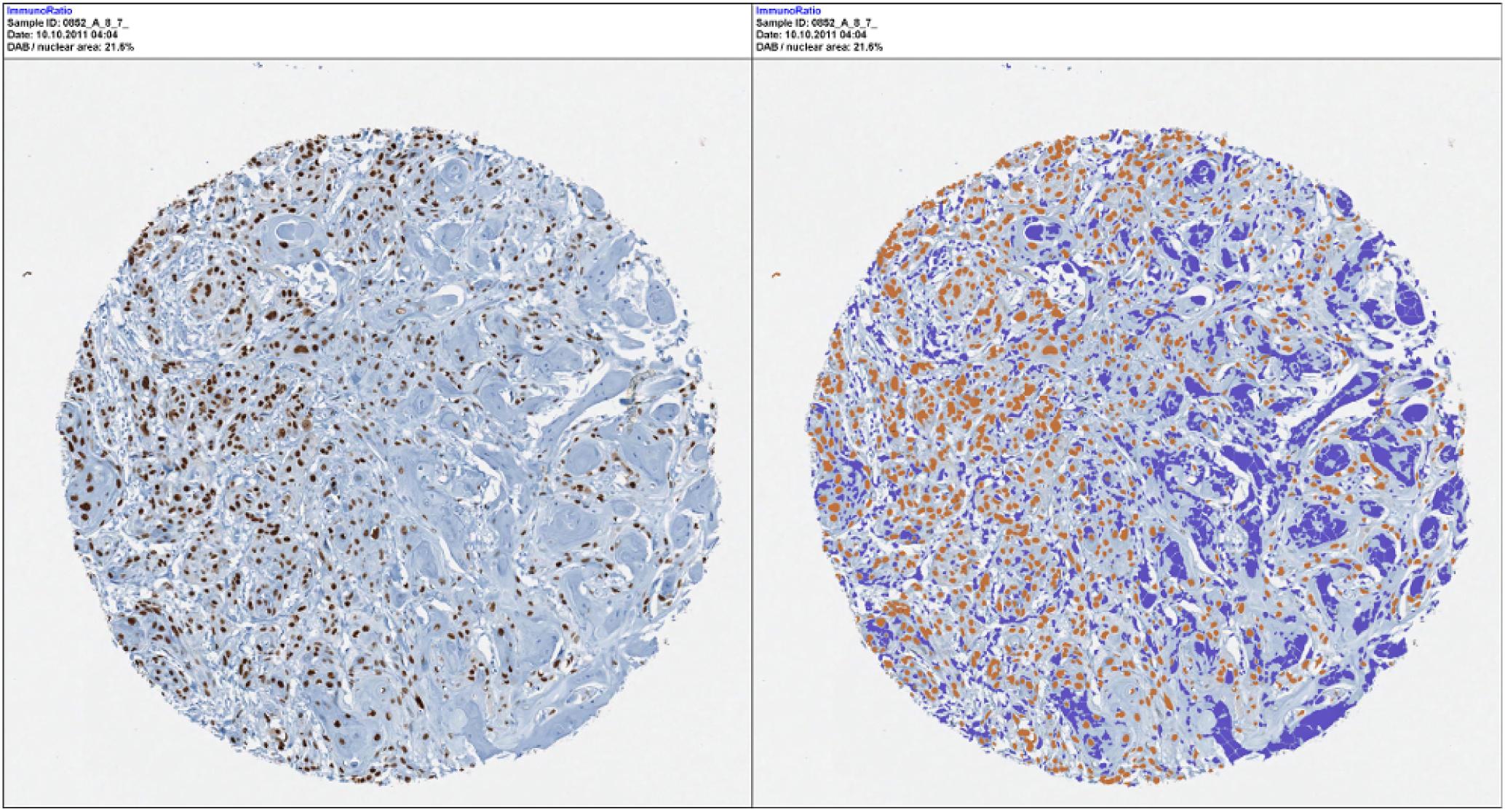
Output of digital analysis for p53 expression. The left figure is the original p53-stained tissue microarray spot. The right figure shows the results of the digital analysis. Detailed description of the algorithm used for digital analysis is provided at http://imtmicroscope.uta.fi/immunoratio.

Percentages of p53 immunohistochemical expression were higher with the visual method (mean 23.4%, SD 32.4%, range from 0% to 99%) than with the digital method (mean 4.7%, SD 7.6%, range from 0% to 43.4%). This difference was statistically significant (P = 8.7*e* − 20).

Correlation between the visual and the digital methods was strong and statistically significant (Spearman’s *ρ* = 0.76, *ρ* = 2.4*e* − 28, Figure 2). Expression levels of p53 according to histologic grade and histologic subtype by visual and digital estimations are shown in Table 1.

**Figure 2:**
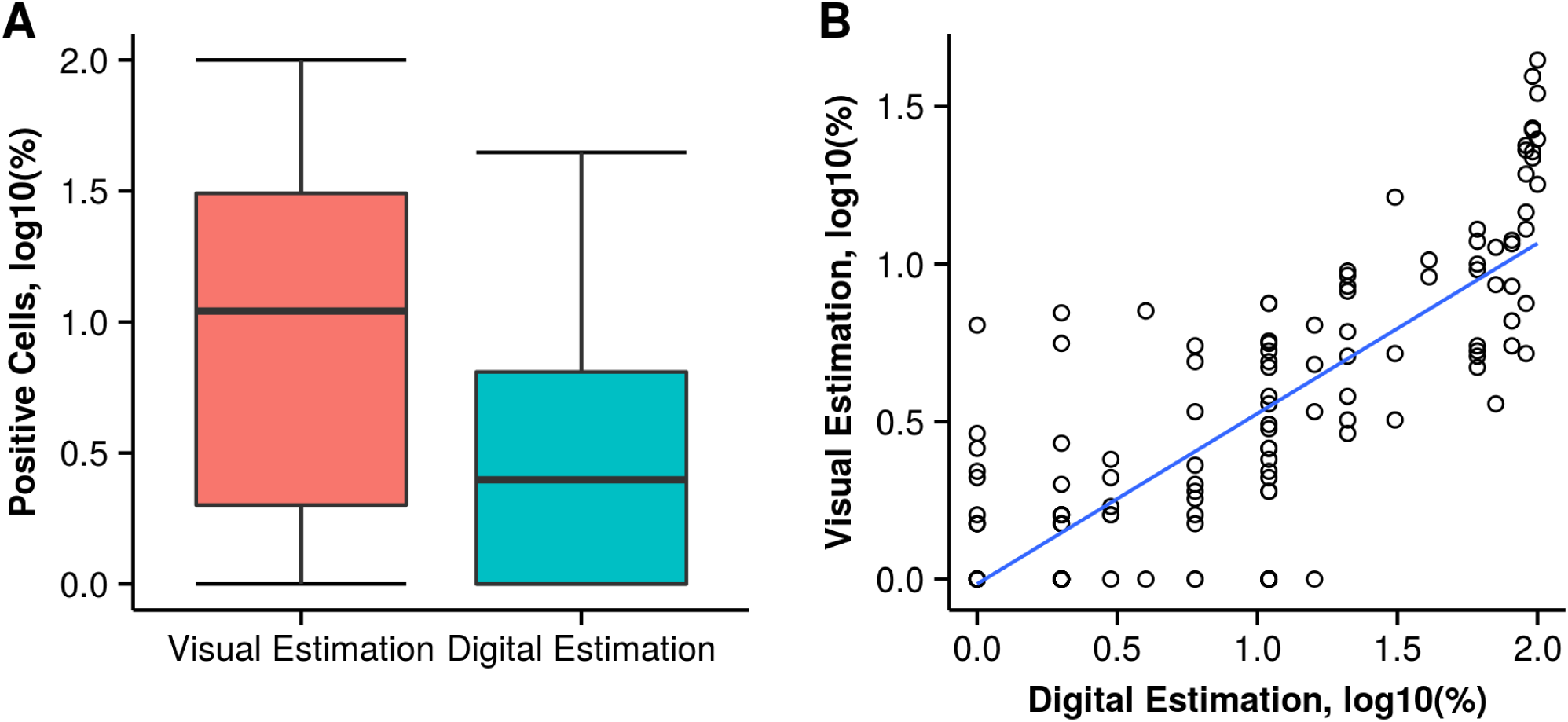
Immunohistochemical Expression of p53 by Visual and Digital Evaluation. A) Boxplots showing the distribution of p53 positive cells with higher percentages using the visual estimation; B) Scatter plot of p53 immunohistochemical expression by visual estimation and digital analysis showing a high correlation between these 2 methods. The blue line represents the regression line. For both plots base-10 logarithmic transformation of p53 percentages were used to aid in data visualization.

## Discussion

This study demonstrates the feasibility of performing digital image analysis of p53 immunohistochemical expression using freely-available, open-source software packages. Our data suggest that visual analysis tends to overestimate the percentage of p53 positive cells. However, the high correlation between visual and digital analyses indicates that both methods are appropriate for estimating the relative levels of immunohistochemical expression. As described in this study, digital estimation of p53 expression could offer an inexpensive and more reliable approach for evaluating the results of immunohistochemical techniques. This approach would also be less time-consuming, less prone to interobserver variability, and the printed output could be easily added to the pathology report. The use of standardized methods would also allow the direct comparison of different studies, and the selection of clinically-applicable thresholds for defining diagnosis or treatment.

In normal cells, the protein p53 plays a central role in the regulation of the cell cycle. Additionally, mutations in the tumor suppressor gene TP53, located on chromosome 17p13, have been identified in approximately 70% of adult solid tumors.^6^ Mutation of TP53 leads to either loss of the protein expression or, more frequently, expression of a mutant protein.^3^ This mutant p53 then accumulates, resulting in an overexpression of the protein. This overexpression can be afterwards detected and quantified using standard diagnostic tools such as immunohistochemistry.

Expression levels of p53 have been used as prognostic tools in several malignancies, including genitourinary tumors.^7^ On this regard, several studies have suggested that immunohistochemical p53 expresión is associated with prognosis in penile carcinomas. Lopes *et al* studied 82 patients with penile cancer treated by penectomy and bilateral inguinal lymphadenectomy.^8^ They found that p53 positivity was associated with lymph node metastasis in univariate and multivariate models. They also found that patients with p53 negative tumors had a better 5-year and 10-year overall survival compared to those patients with p53 positive tumors. Martins *et al* studied 50 patients with penile cancer treated by penectomy.^9^ They found that p53 positivity was associated with tumor progression and disease-specific survival in both univariate and multivariate models. Finally, Zhu et al evaluated p53 expression in 73 patients treated by penectomy and inguinal lymphadenectomy.^10^ They found that p53 positivity was associated with lymph node metastasis and cancer-specific survival in univariate and multivariate models. Besides its potential usefulness as prognosticator of outcome in penile carcinomas, p53 positivity could also aid in the differential diagnosis of penile intraepithelial lesions.^11^ In this scenario, the accurate and reproducible evaluation of p53 immunohistochemical expression could have profound clinical implications.

In summary, evaluation of p53 immunohistochemical expression is feasible using open-source software packages for digital image analysis. Although our analysis was limited to penile SCC, the rationale should also hold for other tumor types in which evaluation of p53 expression is required. This approach would reduce interobserver variability, and would provide a standardized method for reporting the results of immunohistochemical stains. As these diagnostic tools are freely available over the Internet, researchers and practicing pathologists could incorporate them in their daily practice without increasing diagnostic costs.

## Acknowledgments

We are in debt to Helen Fedor and Marcela Southerland, from the TMA Lab Core; Rajni Sharma, PhD, from the Immunopathology Lab; and Kristen L. Lecksell, BS, from the Pathology Department, at the Johns Hopkins Medical Institutions (Baltimore, MD).

